# Genomic characterisation of an *mcr-1* and *mcr*-3-producing *Escherichia coli* strain isolated from pigs in France

**DOI:** 10.1101/2021.09.02.458809

**Authors:** Afaf Hamame, Bernard Davoust, Jean-Marc Rolain, Seydina M. Diene

## Abstract

The current study is about genomic characterisation of an atypical multidrug-resistant *Escherichia coli* harbouring two mobilised colistin resistance (*mcr*) genes isolated from pigs in France. Stool samples taken from a pig farm in Avignon in the department of the Vaucluse were subjected to a molecular screening for the detection of *mcr* gene variants. These samples were cultured on selective LBJMR medium. Growing bacteria were identified using MALDI-TOF, followed by antibiotic susceptibility testing (AST). Whole genome sequencing (WGS) and bioinformatic genome analysis was performed. The selective culture of stools revealed the presence of an *E. coli* strain named Q4552 which was simultaneously positive for the *mcr-*1.1 and *mcr-*3.5 genes. This strain exhibited resistance phenotype to fourteen antibiotics, including colistin. Genome sequencing revealed a circular chromosome and eight plasmids. Genomic analysis revealed a chromosomic integration of a mobile genetic element (MGE) harbouring the *mcr*-1.1 gene, while the *mcr*-3.5 gene was plasmidic (i.e., an IncFII plasmid). Its resistome exhibited twenty-two resistance genes, explaining its multidrug resistance phenotype. The Q4552 strain is an ST-843 clone belonging to the clonal complex Cplx-568 and is the only ST type of this cplx-568 which has been isolated from animals, humans, and the environment. Here, we report the first co-occurrence of the *mcr*-1 and *mcr*-3 genes in France from a pathogenic *E. coli* strain isolated from a pig farm. Since this clone (ST-843) has been reported in zoonotic transmissions, programs to monitor such colistin resistant bacterium are urgently required to avoid its spread and zoonotic transmission to humans.

## INTRODUCTION

Colistin (polymyxin E) is a polycationic antibiotic that interacts with the lipid A of lipopolysaccharide (LPS) in the outer membrane of Gram-negative bacteria (GNB) leading to cell lysis (1). This antibiotic is considered as one of the last-line antimicrobial agents to treat infections caused by multidrug resistant (MDR) Gram-negative bacteria, especially *Enterobacteriaceae* such as *Escherichia coli* and *Klebsiella pneumoniae* carbapenemases-producers (2). Furthermore, the use of polymyxins in veterinary medicine has become very common within the European Union and European Economic Area (EU/EEA).(3) The widespread use of this antibiotic has caused a significant increase in colistin resistance, especially in *E. coli* from pigs. Consequently, as reported, the emergence of colistin resistant bacteria in animals and humans is higher than before (4). Previously, it has been shown that colistin resistance is mediated by chromosomal gene mutations leading to amino acid substitutions in protein sequences associated with this resistance (5–7). Gram-negative bacteria, especially *Enterobacteriaceae*, have developed several mechanisms against colistin including amino acid substitutions in the two-component system *PhoP*/*PhoQ* and *PmrA*/*PmrB*, leading to the addition of phosphoethanolamine and 4-amino-4-deoxy-L-arabinose to the lipid A (8). Other mechanisms, such as the inactivation of the *phoP*/*phoQ* negative feedback regulator (*mgrB*) or overexpression efflux pumps have also been reported (9). Liu *et al*. reported the first *E. coli* harbouring the plasmid bearing mobile colistin resistance *mcr-*1 gene, which they isolated from pigs in China (10). After this, ten other *mcr* gene variants were discovered (11–18). Almost all of these variants are abundant in pigs and poultry farming due to the use of polymyxin to prevent infections caused by bacterial pathogens (19).

In this study, we report the first co-occurrence of two mobile colistin resistance genes, *mcr-*1, and *mcr-*3, in an *E. coli* Q4552 strain isolated from pigs in France.

## RESULTS

### Gene screening, culture, and bacterial identification from stool samples

the RT-PCR screening on extracted DNA from the collected stool samples revealed one positive result. This sample was positive for the *mcr-*1 and *mcr-*3 genes, with CT values of 24 and 21, respectively. According to this result, the selective culture of this sample on the LBJMR medium gave pure bacterial colonies after 24 hours of incubation, then was identified using MALDI-TOF as *E. coli*, with an identification score of 2.4, and was named the Q4552 strain. As shown in **Figures 1A** and **1B**, the *E. coli* Q4552 strain was resistant to colistin with MIC_COL_= 4µg/ml. Moreover, the standard PCR performed on this Q4552 strain and sequencing confirmed the co-presence of the *mcr*-1.1 and *mcr*-3.5 gene variants (**Fig. 1C**). The AST results revealed that the *E. coli* Q4552 strain was resistant to different β-lactams, including amoxicillin, amoxicillin/clavulanic acid, and penicillin G, but remained susceptible to cephalosporins and carbapenems (**Table 1**). Overall, this Q4552 strain was resistant to fourteen different antibiotics tested (**Table 1**).

**Table 1:** Phenotypic and genotypic comparison of *E. coli* Q4552 according to antibiotic family

| Antibiotic Family | Antibiotics | CMI (mm) | CMI (mg/ml) | ARGs | encoded gene for | % COVERAGE ARG ANNOT | % COVERAGE Resfinder |
| --- | --- | --- | --- | --- | --- | --- | --- |
| **Penicillins** | Penicillin A: Amoxicillin | R:0 | | (Bla)AmpC2 | β-lactamase Class C | 99% | 100% |
| | Amoxicillin + Clavulanic acid | R:16 | >256 | (Bla)AmpC1 | β-lactamase Class C | 99.88% | 100% |
|  | Oxacillin | R | >256 | (Bla)blaTEM | BLSE Class A | 100% | 100% |
|  | Piperacillin | S | 2 |  |  | 100% | 100% |
|  | Aztreonam | S | 0.46 |  |  |  |  |
|  | Penicillin G | R | >32 |  | Carboxypeptidase endopeptidase |  |  |
|  | Tazobactam piperacillin | S: 24.2 | 0.38 |  |  |  |  |
| **Cephalosporins** | Cephalosporin G1: cefalotin | I:16,1 | 0.32 | acrF | Inner membrane transporter | 100% |  |
|  | Cephalosporin G2: cefotaxime | S | <0.25 | AcrE | Membrane fusion protein | 99.66 % |  |
|  | Cephalosporin G3: ceftriaxone | S:38,8 |  | acrS | Efflux pump | 99% |  |
|  | Cephalosporin G4: cefepime | S:30 |  |  |  |  |  |
| **Carbapenems** | Meropenem | S | <0.125 |  |  |  |  |
|  | Ertapenem | S: 39.3 | 0.03 |  |  |  |  |
|  | Imipenem | S:35.6 | <0.32 |  |  |  |  |
| **Macrolides** | Clindamycin | R | >256 | erm(B)_1 |  | 100% | 100% |
|  | Erythromycin | R | >256 | mph(A) |  | 100% | 100% |
|  | Levofloxacin | S | 0.125 | emrA, emrB, emrR | Multidrug efflux pump | 99% |  |
| **Quinolone** | Ciprofloxacin | R | 1,024 | mdtH | Multidrug resistance protein | 97.87% |  |
| **Tetracycline** | Doxycycline | R:10.4 | 32 | (Tet)tetA, | Tetracycline efflux pump | 94.12% | 99,92 |
|  | Minocycline |  | 12 | (Tet)tetR | Tetracycline efflux pump | 100% |  |
| **Sulfamids** | Trimethoprim sulfamethoxazole | R:0 |  | dfrA12_8 |  |  | 100% |
|  | Sulfamide | R | 64 | (Sul)sul3 |  | 100% | 100% |
| **Aminoglycosides** | Streptomycin | R |  | (AGly)aadA1-pm<br>(AGly)aadA2 | Aminoglycoside 6-Acetyltransferase | 93.86%<br>100% | 98.48%<br>95.24% |
|  | Gentamicin | S:37.6 | <0.64 |  |  |  |  |
|  | Netilmicin |  | 0.25 |  |  |  |  |
|  | Kanamycin |  | 1.5 |  |  |  |  |
|  | TGC Tigecycline |  | 0.5 |  |  |  |  |
|  | Amikacin | S:25.6 | 1.5 |  |  | 100% |  |
| **Phenicol** | chloramphenicol |  |  | (Phe)cmlA1 | Mutation | 100% |  |
| **Polymyxin E** | Colistin | R: 13,6 | 4 | (Col) mcr-1.1 | Phosphatidylethanolamine<br>Transferase | 100% | 100% |
|  |  |  |  | (Col) mcr-3.5 |  | 100% | 100% |
| **OTHERS** | Fosfomycin | R | 0.125 | mdtG | Multidrug efflux pump | 99.27 % |  |
|  | Metronidazole | R | >256 |  |  |  |  |
|  | Rifampicin | R | >32 |  |  |  |  |

**Figure 1:**
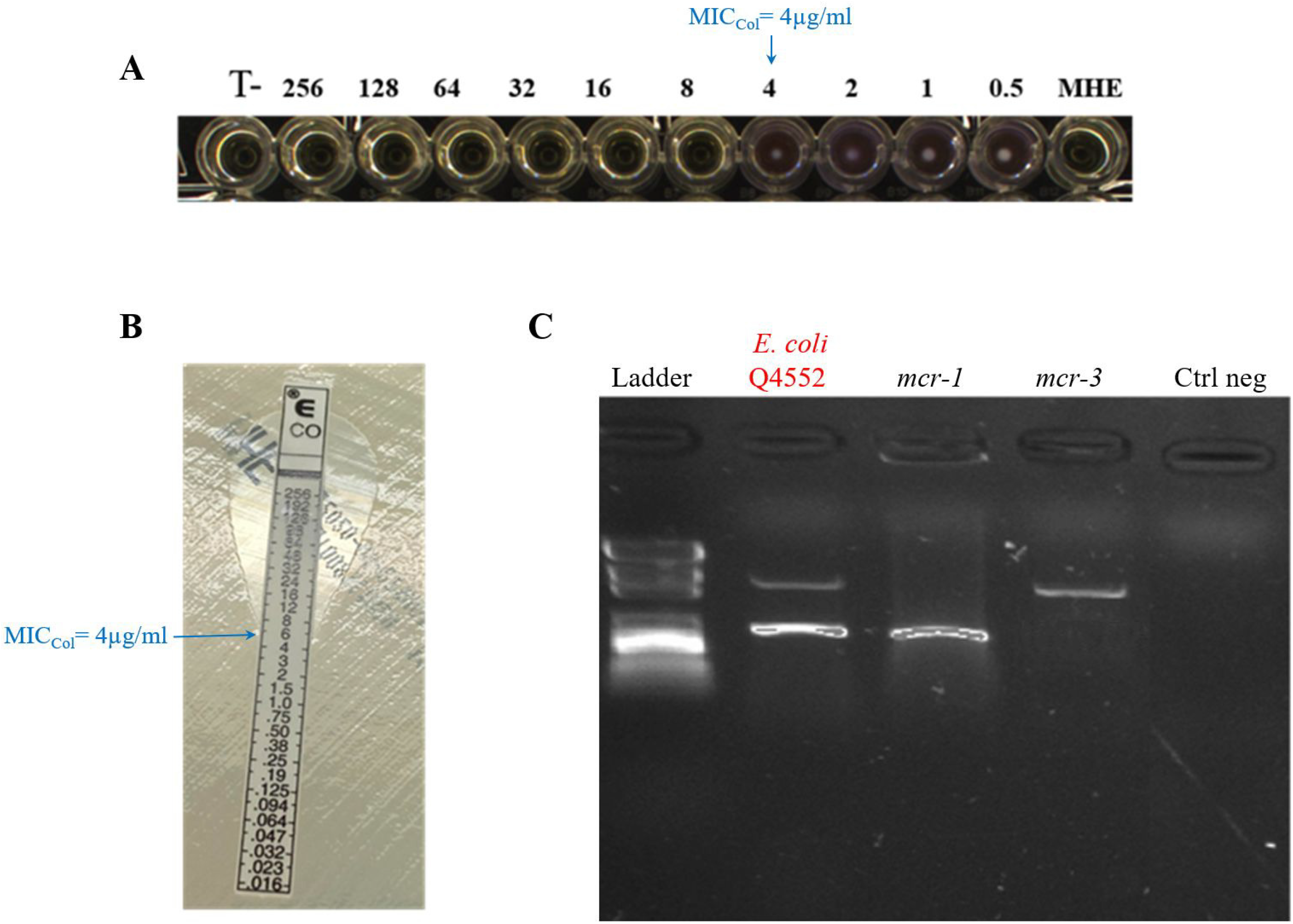
Phenotypic characterisation of the *E. coli* Q4552 strain carrying the two *mcr* variants. **A:** colistin susceptibility test according to EUCAST Microdilution UMIC. **B:** Colistin E-test on *E. coli* Q4552 strain. **C:** Multiplex standard PCR targeting both *mcr*-1 and *mcr*-3.

### Genomic analysis of the colistin resistant Q4552 strain

genome assembly using combined raw reads from the Illumina and Nanopore technologies resulted in the circular chromosome with 4,778,049 bps with 80.87% GC content and eight circular plasmids with sizes which ranged from 84,331 bps to 3,607 bps and a %GC content ranging from 59.94% to 43.39% (**Table 2**). As presented in **Table 1**, resistome analysis reveals the presence in the genome of several genes encoding for resistance to β-lactams (namely *bla*_AMPC2_, *bla*_AMPC1_, *bla*_TEM_, and *bla*_ampH_), to aminoglycosides, especially streptomycin (*aadA*1 and *aadA*2), to macrolides, especially clindamycin and erythromycin (*ermB* and *mphA*), and to quinolones (*mdtH*).

**Table 2:**
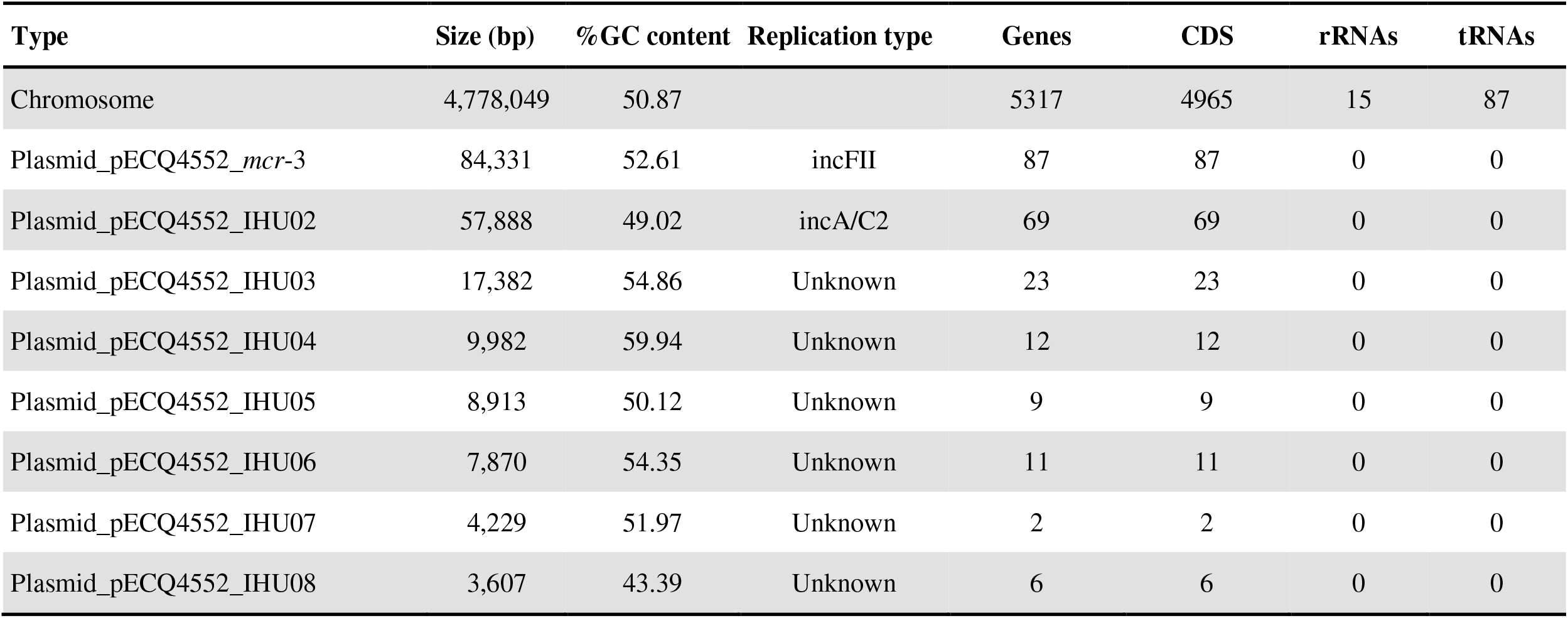
Genomic features of the *E. coli* Q4552 strain

Resistance to colistin was mediated by the presence of the *mcr-*1.1 and *mcr-*3.5 genes. Interestingly, sequence analysis shown that the *mcr*-1.1 gene was located a mobile genomic element (MGE) with 51,089 bps from the Q4552 chromosome. As shown in **Figure 2**, the *E. coli* Q4552 chromosome was compared with published sequences, including the chromosome sequence of *E. coli* K-12 MG1655, *E. coli* CP66_6, and *E. coli* plasmid pPN43. Our analysis reveals that the MGE: *mcr*-1.1 was integrated within a specific integration site upstream of a tRNA gene and mediated by a phage integrase *intA*, as compared to the *E. coli* K-12 MG1655 chromosome (**Fig. 2B**). Moreover, within this MGE, the *mcr*-1 gene is flanked on both sides by the IS30, which is suggestive of a transposition event within this MGE. While this MGE is also identified within the chromosome of the *E. coli* CP66_6 strain isolated from a pig in Sichuan (China), this MGE is, furthermore, identified within the published plasmid *E. coli* pPN43 strain from a human sample in Thailand (**Fig. 2B**). Interestingly, in addition to the *mcr*-1 gene, a restriction-modification system type 1 (RMS type-1) and toxin/antitoxin system were identified within this MGE (**Fig. 2B**) suggestive of a stabilization of MGE within the genome. Regarding *mcr*-3.5, our genomic analysis revealed that this was located in one of the eight identified plasmids (pQ4552_*mcr*-3, an IncFII type) (**Fig. 3**). As shown in **Figure 3**, this plasmid pQ4552_*mcr*-3 (84,331 bps; 96.56%CG) was compared with the six most similar plasmids from the NCBI database, with an alignment percentage between 78% and 98% and an identity percentage between 98.60 and 99.96%. This analysis shows that *mcr*-3 is only present in two out of the six plasmids, including the *E. coli* plasmid pVE796 (AP018353.1) and the unnamed *K. pneumoniae* plasmid (CP041102.1). A toxin/antitoxin (*pemI*/*pemK*) system allows this plasmid to be maintained within this *E. coli* Q4552 strain (**Fig. 3**).

**Figure 2:**
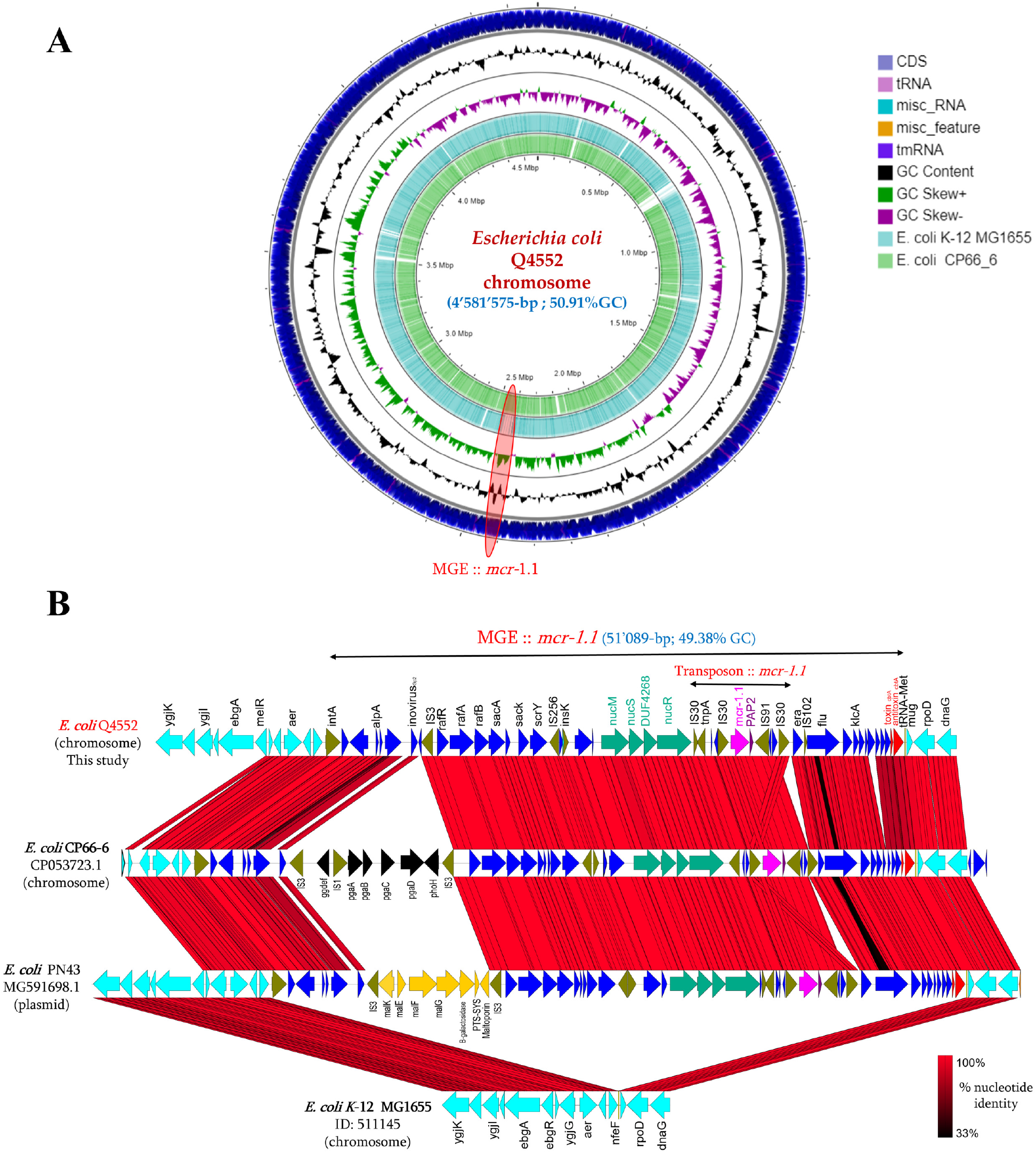
**A**. Circular maps of the chromosomal genome with *mcr-1*. **B**. Zoom on the genetic environment of the transposon carrying the *mcr-1*.*1* gene compared with the top best hit blast *E. coli* CP66-6 and plasmid *E. coli* PN43.

**Figure 3:**
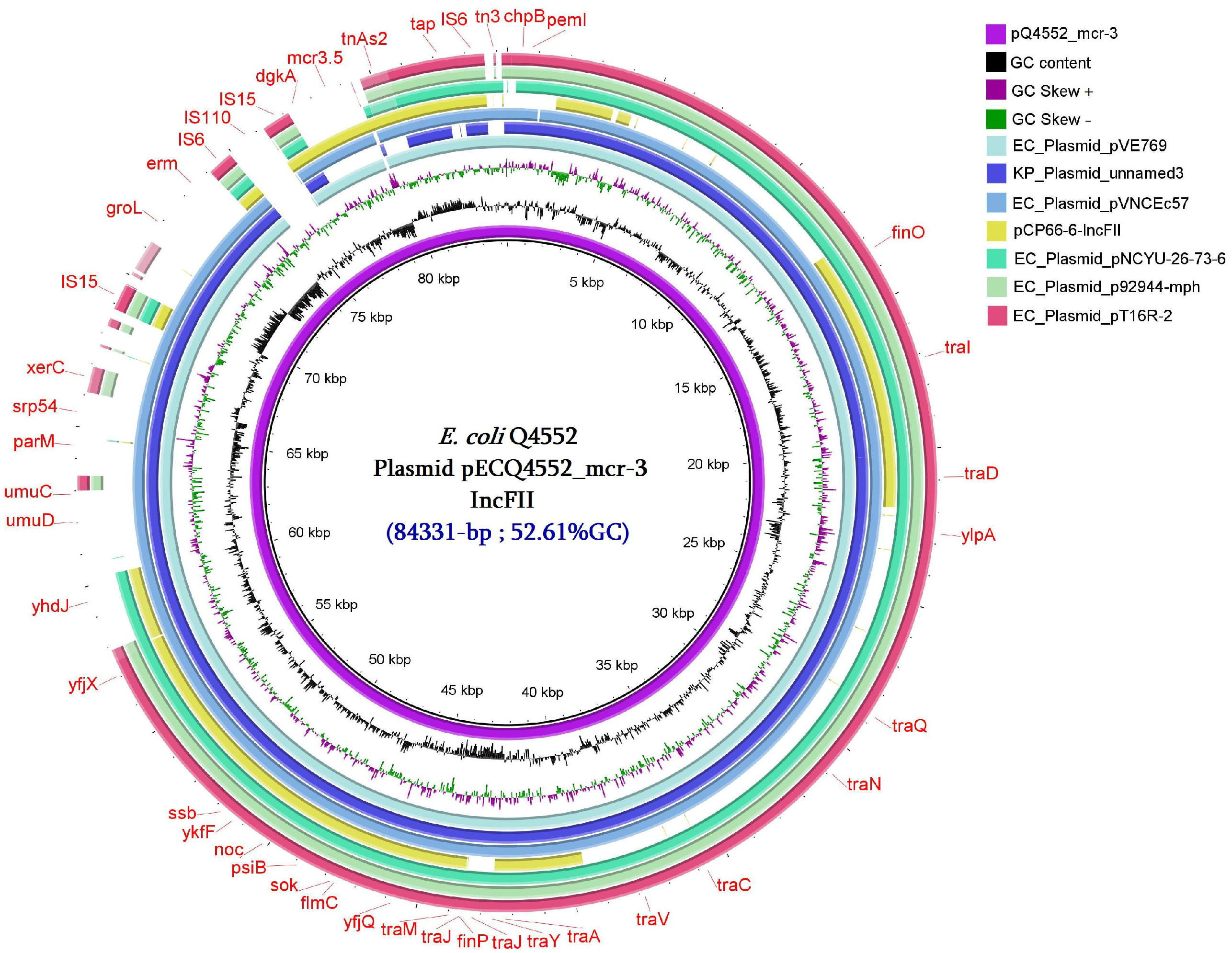
Circular map of the plasmid carrying *mcr-3*.*5* compared to six homologous plasmids with considerable query coverage.

### Multilocus sequence typing analysis

to investigate the clonality of the Q4552 strain, an MLST analysis was performed from the genome sequence, which classified the colistin-resistant Q4552 strain into ST-843, belonging to the clonal complex Cplx-568. As presented in **Figure 4**, the Cplx-568 complex is thus far composed of 14 ST types from the Institut Pasteur database. The investigation of the isolation sources of these ST types revealed diverse sources, including humans, animals, and the environment. Interestingly, three ST types, namely ST-80, ST-1946, and ST-843, were isolated from humans and animals, suggesting the ability of this clone to be transferred from animals to humans and vice versa. However, only the ST-843 (our Q4552) was isolated from the three different sources (animal, human, and the environment) (**Fig. 4**).

**Figure 4:**
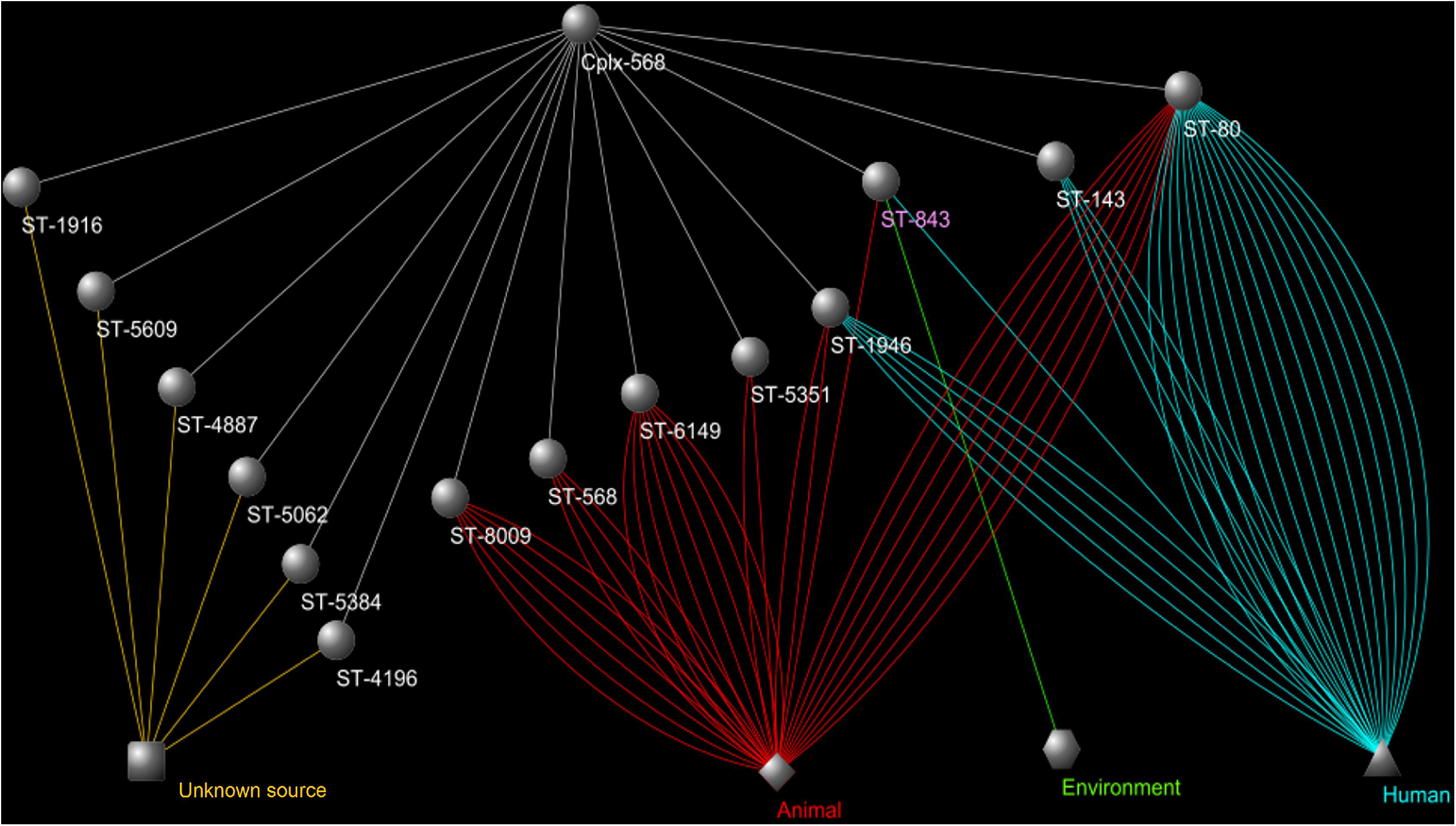
MLST and cytoscape analysis of *E. coli* Q4552 (ST843) clone compared with the other clones associated with Ccpx568.

## DISCUSSION

According to the literature, the main source of the strains with a co-occurrence of *mcr-*1 and *mcr-*3 are animals. Here, the ecosystem of *E. coli* Q4552 is also farm animals, since this strain was isolated from a pig sample. Pigs are in close contact with humans. Depending on culture and traditions, pig meat is consumed around the entire world in various forms, and humans are consequently exposed to contamination by strains with the *mcr* genes, as already reported in farm workers and the surrounding environment being largely contaminated by the *mcr*-1 gene (20). So far, few studies have reported the co-occurrence of two *mcr* genes within a bacterium. This co-occurrence has been reported in an *E. coli* strain isolated from cattle (21) in Spain, pigs in Thailand (22), and in dog faeces from China (23). Moreover, the co-occurrence of *mcr*-1 and *mcr*-3 has also been reported in humans, such as the *E. coli* strain isolated from a human urine sample in New Zealand (24), and a *K. pneumonaie* strain isolated from a stool sample from a healthy volunteer in Thailand (25). The identified MGE harbouring the *mcr*-1.1 gene from the Q4552 chromosome appears to be extremely rare in available genomes from the NCBI database, and has only been detected in an *E. coli* CP66-6 strain with *mcr*-1 which was isolated, as was ours, from a pig in China (26), but also from a recombinant plasmid of an *E. coli* strain isolated from a human in Thailand (27). However, both *E. coli* strains are not ST-843 like our strain, but share this MGE. This MGE seems to be stabilised on the *E. coli* Q4552 chromosome given the presence of RMS type 1 and the toxin/antitoxin system within this acquired MGE. While the *mcr*-1 gene was identified from the Q4552 chromosome, the *mcr*-3.5 gene variant was identified on a conjugative and recombinant IncFII type plasmid. Highly similar plasmids harbouring the *mcr*-3.5 gene have been reported in the literature from *E. coli* strains isolated in Vietnam (AP018353.1), Thailand (Cp041102.1) and China (26). *E. coli* Q4552 is an atypical bacterium which belongs to the clonal complex Ccpx-568 and has an ST-843. ST-843 appears closely related to ST-80 and ST-1946 which are two *E. coli* clones well reported in zoonotic transmissions from animals and humans. Indeed, ST-80 has well been reported as a uropathogenic *E. coli* clone from retail food such as chicken and is able to cause community acquired infections (28).

In conclusion, *E. coli* Q4552 exhibited a multidrug resistance phenotype mediated by various resistance genes. The most unusual co-occurrence is the fact that *mcr-*1.1 is chromosomic, with more than 4 IS transposases and a plasmidic *mcr-*3.5. As reported in the literature, *mcr* genes encode for phosphoethanolamine transferase proteins and could serve as defensive enzymes against phage attacks (29). These results give rise to questions about the real functions of these *mcr* genes within this Q4552 strain, which was not more resistant to colistin, despite this co-occurrence.

## MATERIALS AND METHODS

### Decree and law authorising the use of animal products for research and diagnosis

Prefectorial authorisation (Bouches-du-Rhône) No.13 205 107 of 4 September 2014 authorising the IHU Méditerranée Infection to use unprocessed animal by-products in categories 1, 2 and 3 for research and diagnostic purposes. Order of 8 December 2011 laying down health rules concerning animal by-products and derived products in application of Regulation (EC) No. 1069/2009 and Regulation (EU) No. 142/2011. In 2020, pig faecal samples were collected from a pig fattening farm near Avignon (43° 50’ 32.5’’N - 4° 57’ 33.9’’E), in the department of the Vaucluse, France. The pigs are crossbreeds (mother: Large White X Landrace cross - father: Pietrain X Duroc cross). These animals (2 to 8 months old) were sampled in several batches. After fattening, they are intended for human consumption in Corsica, France.

### DNA extraction

DNA was extracted from the pig faecal samples according to a protocol used in previous studies (30). 1g of faeces, Buffer G2 and proteinase K were incubated at 56°C overnight and extracted using the EZ1 DNeasy Blood Tissue Kit (Qiagen GmbH, Hilden, Germany) in line with the manufacturer’s protocol.

### Molecular screening of colistin resistance genes

the presence of genes encoding for colistin resistance (*mcr*-1, *mcr-*2, *mcr*-3, *mcr*-4, *mcr*-5, *mcr*-8) was investigated by Real-Time PCR assay(31),(32) using the CFX96 TM Real time System / C1000 TM Touch thermal Cycler (Bio-Rad, Singapore). Targeted genes were detected using specific primers and probes (**Suppl. Table S1**) (33). Results were considered positive when the cycle threshold value of real-time PCR was ≤30. Standard PCR multiplex was then performed to confirm positive qPCR results, followed by sequencing.

### Screening of colistin resistant Gram-negative bacteria

the selected faecal sample was enriched in TSB for 72 hours then cultured on a selective medium known as LBJMR (Lucie Bardet Jean Marc Rolain), containing colistin sulfate salt at a concentration of 4µg/ml, and vancomycin (50µg/ml), with glucose as the fermentative substrate, on a Purple Agar Base. Using this LBJMR medium, *Enterobacteriaceae* isolates are yellow in colour, contrasting with the purple agar and are of a different size (2–3 mm), whereas *Enterococci* isolates are small and round (0.1–1 mm) (34).

### MALDI-TOF MS identification

colonies with different morphologies were picked up from the LBJMR agar plate. Those colonies were spotted and identified using a Microflex LS spectrometer (Bruker Daltonics, Bremen, Germany) (35). The strain was properly identified at the species level when the score is ≤ 2.

### Antibiotic susceptibility testing (AST)

the Disk diffusion (DD) test (Kirby-Bauer procedure) is a routine susceptibility testing method which was implemented according to the CLSI and EUCAST guidelines (https://www.eucast.org/clinical_breakpoints/). Sixteen antibiotics were tested, including ampicillin, amoxicillin-clavulanic acid, ceftazidime, cefotaxime, cefepime, cefoxitin, piperacillin-tazobactam, aztreonam, meropenem, ertapenem, imipenem, tigecycline, ciprofloxacin, gentamicin, trimethoprim-sulfamethoxazole, and colistin (Bio-Rad, Marne-la-Coquette, France). Additional AST methods such as the E-test (BioMérieux) and microdilution using UMIC microdilution (Biocentric, Bandol, France) were performed to confirm some AST results, as previously reported.

### Genome sequencing and bioinformatic analysis

whole genome sequencing was performed using MiSeq Illumina (36) (Illumina Inc., San Diego, CA, United states) and Nanopore MINION technologies (37). Genome assembly was performed using a Canu assembler, which combines Illumina and Nanopore sequence reads, and genome annotation was performed using Prokka.(38) Resistome analysis was investigated using different databases including Resfinder (39), ARG-ANNOT (40), Card (41), and Plasmid Finder (42). CGview software was then used for circular representation of chromosome and plasmid sequences (43).

### Multilocus Sequence Typing (MLST)

sequence type analysis was carried out from genomic sequences and the set of close typing sequences was determined by PubMLST then compared with the 11,808 ST of *Escherichia* spp (https://pubmlst.org/) using Cytoscape and layout software (44).

### Sequence accession number

Complete genome sequence including the chromosome and the eight plasmids of the *E. coli* Q4552 strain have been deposited in the NCBI GenBank database under accession numbers: CP077063, CP077064, CP077065, CP077066, CP077067, CP077068, CP077069, CP077070, and CP077071.

## Funding information

This work was supported by the French Government under the “Investissements d’avenir” (Investments for the Future) programme managed by the Agence Nationale de la Recherche (ANR, fr: National Agency for Research), (reference: Méditerranée Infection 10-IAHU-03).

## Conflicts of interest

The authors declare that they have no competing interests.

## Authors’ contributions

SMD, BD, and JMR designed the study; AF performed microbiology analyses; AF and SMD drafted the manuscript; AF, BD, JMR, and SMD made corrections and critical revisions. All the authors have read and approved the final version of the manuscript.

---

**Supplementary Table S1:**
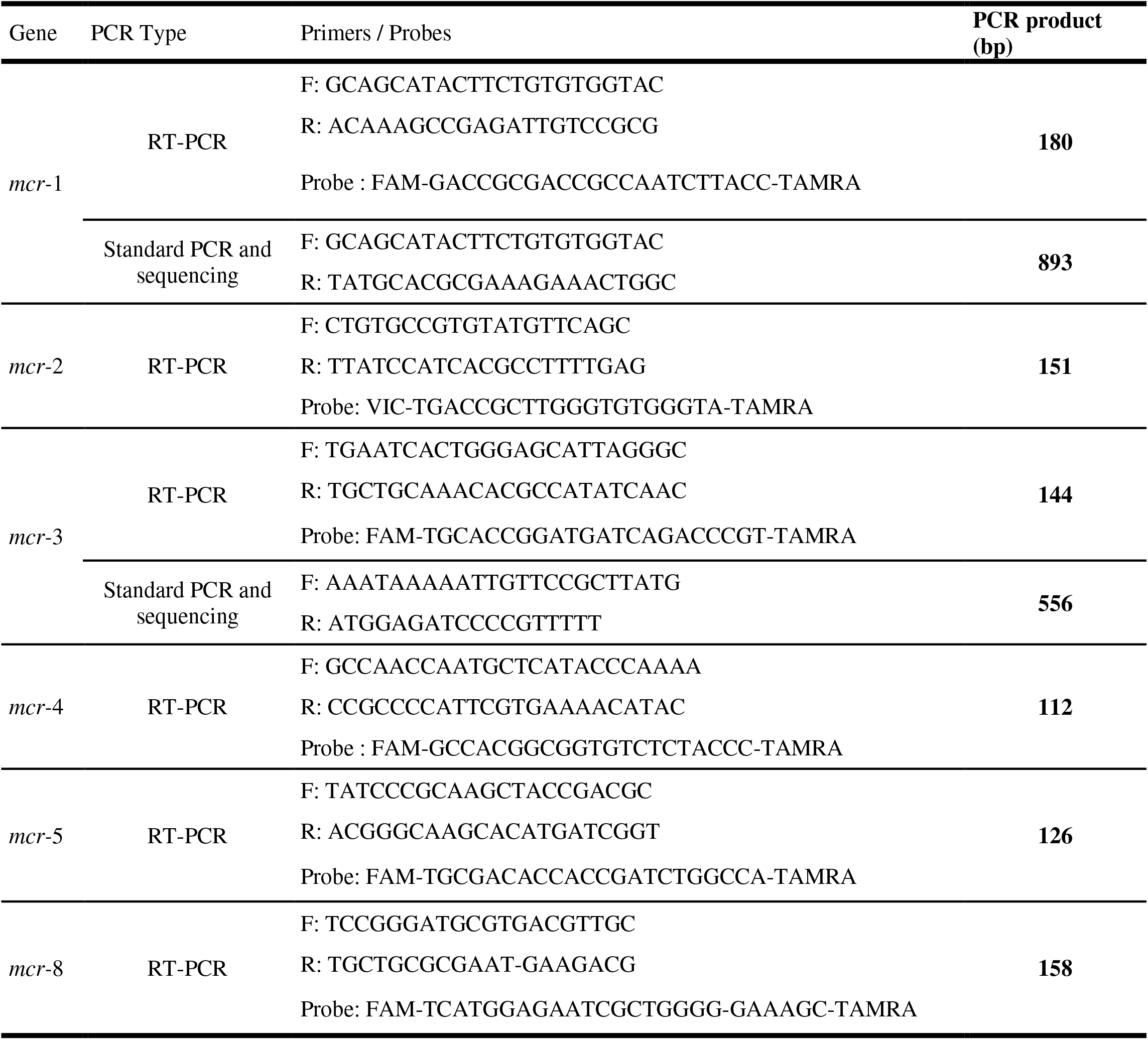
Sequences of primers and probes used for real-time PCRs and conventional PCRs in this study (45, 46).

## REFERENCES

1. Hamel M, Rolain JM, Baron SA. 2021. The history of colistin resistance mechanisms in bacteria: Progress and challenges. Microorganisms 9:1–18.

2. Poirel L, Jayol A, Nordmanna P. 2017. Polymyxins: Antibacterial activity, susceptibility testing, and resistance mechanisms encoded by plasmids or chromosomes. Clin Microbiol Rev. American Society for Microbiology.

3. Catry B, Cavaleri M, Baptiste K, Grave K, Grein K, Holm A, Jukes H, Liebana E, Navas AL, Mackay D, Magiorakos AP, Romo MAM, Moulin G, Madero CM, Pomba MCMF, Powell M, Pyörälä S, Rantala M, Ružauskas M, Sanders P, Teale C, Threlfall EJ, Törneke K, Van Duijkeren E, Edo JT. 2015. Use of colistin-containing products within the European Union and European Economic Area (EU/EEA): development of resistance in animals and possible impact on human and animal health. Int J Antimicrob Agents. Elsevier B.V.

4. Hadjadj L, Riziki T, Zhu Y, Li J, Diene SM, Rolain JM. 2017. Study of mcr-1 gene-mediated colistin resistance in enterobacteriaceae isolated from humans and animals in different countries. Genes (Basel) 8:1–16.

5. Johansen HK, Moskowitz SM, Ciofu O, Pressler T, Høiby N. 2008. Spread of colistin resistant non-mucoid Pseudomonas aeruginosa among chronically infected Danish cystic fibrosis patients. J Cyst Fibros 7:391–7.

6. Arduino SM, Quiroga MP, Ramírez MS, Merkier AK, Errecalde L, Di Martino A, Smayevsky J, Kaufman S, Centrón D. 2012. Transposons and integrons in colistin-resistant clones of Klebsiella pneumoniae and Acinetobacter baumannii with epidemic or sporadic behaviour. J Med Microbiol 61:1417–1420.

7. Tenover FC, McGowan JE. 1996. Reasons for the Emergence of Antibiotic Resistance. Am J Med Sci 311:9–16.

8. Baron S, Hadjadj L, Rolain JM, Olaitan AO. 2016. Molecular mechanisms of polymyxin resistance: knowns and unknowns. Int J Antimicrob Agents 48:583–591.

9. Olaitan AO, Morand S, Rolain JM. 2014. Mechanisms of polymyxin resistance: Acquired and intrinsic resistance in bacteria. Front Microbiol 5:1–18.

10. Liu Y-Y, Wang Y, Walsh TR, Yi L-X, Zhang R, Spencer J, Doi Y, Tian G, Dong B, Huang X, Yu L-F, Gu D, Ren H, Chen X, Lv L, He D, Zhou H, Liang Z, Liu J-H, Shen J. 2016. Emergence of plasmid-mediated colistin resistance mechanism MCR-1 in animals and human beings in China: a microbiological and molecular biological study. Lancet Infect Dis 16:161–8.

11. Xavier BB, Lammens C, Ruhal R, Malhotra-Kumar S, Butaye P, Goossens H, Malhotra-Kumar S. 2016. Identification of a novel plasmid-mediated colistinresistance gene, mcr-2, in Escherichia coli, Belgium, june 2016. Eurosurveillance 21:30280.

12. Yin W, Li H, Shen Y, Liu Z, Wang S, Shen Z, Zhang R, Walsh TR, Shen J, Wang Y. 2017. Novel Plasmid-Mediated Colistin Resistance Gene mcr-3 in Escherichia coli. MBio 8.

13. Carattoli A, Villa L, Feudi C, Curcio L, Orsini S, Luppi A, Pezzotti G, Magistrali CF. 2017. Novel plasmid-mediated colistin resistance mcr-4 gene in Salmonella and Escherichia coli, Italy 2013, Spain and Belgium, 2015 to 2016. Euro Surveill Bull Eur sur les Mal Transm = Eur Commun Dis Bull 22.

14. Borowiak M, Fischer J, Hammerl JA, Hendriksen RS, Szabo I, Malorny B. 2017. Identification of a novel transposon-associated phosphoethanolamine transferase gene, mcr-5, conferring colistin resistance in d-tartrate fermenting Salmonella enterica subsp. enterica serovar Paratyphi B. J Antimicrob Chemother 72:3317–3324.

15. Teale C, Anjum MF, Kirchner M, Stubberfield EJ, Nunez-Garcia J, Lemma F, Randall LP, AbuOun M, Dormer L, Duggett NA, Crook DW, Smith RP. 2017. mcr-1 and mcr-2 (mcr-6.1) variant genes identified in Moraxella species isolated from pigs in Great Britain from 2014 to 2015. J Antimicrob Chemother 72:2745–2749.

16. Yang Y-Q, Li Y-X, Lei C-W, Zhang A-Y, Wang H-N. 2018. Novel plasmid-mediated colistin resistance gene mcr-7.1 in Klebsiella pneumoniae. J Antimicrob Chemother 73:1791–1795.

17. Carroll LM, Gaballa A, Guldimann C, Sullivan G, Henderson LO, Wiedmann M. 2019. Identification of novel mobilized colistin resistance gene mcr-9 in a multidrug-resistant, colistin-susceptible salmonella enterica serotype typhimurium isolate. MBio 10:1–6.

18. Wang C, Feng Y, Liu L, Wei L, Kang M, Zong Z. 2020. Identification of novel mobile colistin resistance gene mcr-10. Emerg Microbes Infect 9:508–516.

19. Kempf I, Fleury MA, Drider D, Bruneau M, Sanders P, Chauvin C, Madec JY, Jouy E. 2013. What do we know about resistance to colistin in Enterobacteriaceae in avian and pig production in Europe? Int J Antimicrob Agents. Elsevier.

20. Dandachi I, Fayad E, Sleiman A, Daoud Z, Rolain JM. 2020. Dissemination of Multidrug-Resistant and mcr-1 Gram-Negative Bacilli in Broilers, Farm Workers, and the Surrounding Environment in Lebanon. Microb Drug Resist 26:368–377.

21. Hernández M, Iglesias MR, Rodríguez-Lázaro D, Gallardo A, Quijada NM, Miguela-Villoldo P, Campos MJ, Píriz S, López-Orozco G, de Frutos C, Sáez JL, Ugarte-Ruiz M, Domínguez L, Quesada A. 2017. Co-occurrence of colistin-resistance genes mcr-1 and mcr-3 among multidrug-resistant Escherichia coli isolated from cattle, Spain, September 2015. Eurosurveillance 22:30586.

22. Khine NO, Lugsomya K, Kaewgun B, Honhanrob L, Pairojrit P, Jermprasert S, Prapasarakul N. 2020. Multidrug Resistance and Virulence Factors of Escherichia coli Harboring Plasmid-Mediated Colistin Resistance: mcr-1 and mcr-3 Genes in Contracted Pig Farms in Thailand. Front Vet Sci 7:582899.

23. Du C, Feng Y, Wang G, Zhang Z, Hu H, Yu Y, Liu J, Qiu L, Liu H, Guo Z, Huang J, Qiu J. 2020. Co-occurrence of the mcr-1.1 and mcr-3.7 genes in a multidrug-resistant escherichia coli isolate from China. Infect Drug Resist 13:3649–3655.

24. Creighton J, Anderson T, Howard J, Dyet K, Ren X, Freeman J. 2019. Co-occurrence of mcr-1 and mcr-3 genes in a single Escherichia coli in New Zealand. J Antimicrob Chemother. Oxford University Press.

25. Yu Y, Andrey DO, Yang RS, Sands K, Tansawai U, Li M, Portal E, Gales AC, Niumsup PR, Sun J, Liao X, Liu YH, Walsh TR. 2020. A Klebsiella pneumoniae strain co-harbouring mcr-1 and mcr-3 from a human in Thailand. J Antimicrob Chemother 75:2372–2374.

26. Li R, Du P, Zhang P, Li Y, Yang X, Wang Z, Wang J, Bai L. 2021. Comprehensive Genomic Investigation of Coevolution of mcr genes in Escherichia coli Strains via Nanopore Sequencing Glob Challenges 5:2000014.

27. Yang Q, Li M, Spiller OB, Andrey DO, Hinchliffe P, Li H, MacLean C, Niumsup P, Powell L, Pritchard M, Papkou A, Shen Y, Portal E, Sands K, Spencer J, Tansawai U, Thomas D, Wang S, Wang Y, Shen J, Walsh T. 2017. Balancing mcr-1 expression and bacterial survival is a delicate equilibrium between essential cellular defence mechanisms. Nat Commun 8:1–12.

28. Yamaji R, Friedman CR, Rubin J, Suh J, Thys E, McDermott P, Hung-Fan M, Riley LW. 2018. A Population-Based Surveillance Study of Shared Genotypes of Escherichia coli Isolates from Retail Meat and Suspected Cases of Urinary Tract Infections. mSphere 3.

29. Khedher M Ben, Baron SA, Riziki T, Ruimy R, Raoult D, Diene SM, Rolain JM. 2020. Massive analysis of 64,628 bacterial genomes to decipher water reservoir and origin of mobile colistin resistance genes: is there another role for these enzymes? Sci Rep 10:1–10.

30. Ngaiganam EP, Pagnier I, Chaalal W, Leangapichart T, Chabou S, Rolain JM, Diene SM. 2019. Investigation of urban birds as source of β-lactamase-producing Gram-negative bacteria in Marseille city, France. Acta Vet Scand 61:1–7.

31. Edelstein M, Pimkin M, Palagin I, Edelstein I, Stratchounski L. 2003. Prevalence and Molecular Epidemiology of CTX-M Extended-Spectrum β-Lactamase-Producing Escherichia coli and Klebsiella pneumoniae in Russian Hospitals. Antimicrob Agents Chemother 47:3724–3732.

32. Chabou S, Leangapichart T, Okdah L, Le Page S, Hadjadj L, Rolain JM. 2016. Real-time quantitative PCR assay with Taqman®probe for rapid detection of MCR-1 plasmid-mediated colistin resistance. New Microbes New Infect. Elsevier Ltd.

33. Leangapichart T, Dia NM, Olaitan AO, Gautret P, Brouqui P, Rolain JM. 2016. Acquisition of extended-spectrum β-lactamases by Escherichia coli and Klebsiella pneumoniae in gut microbiota of pilgrims during the hajj pilgrimage of 2013. Antimicrob Agents Chemother 60:3222–3226.

34. Bardet L, Le Page S, Leangapichart T, Rolain JM. 2017. LBJMR medium: A new polyvalent culture medium for isolating and selecting vancomycin and colistin-resistant bacteria. BMC Microbiol 17:1–10.

35. Singhal N, Kumar M, Kanaujia PK, Virdi JS. 2015. MALDI-TOF mass spectrometry: An emerging technology for microbial identification and diagnosis. Front Microbiol. Frontiers Research Foundation.

36. Caporaso JG, Lauber CL, Walters W a, Berg-Lyons D, Huntley J, Fierer N, Owens SM, Betley J, Fraser L, Bauer M, Gormley N, Gilbert J a, Smith G, Knight R. 2012. Ultra-high-throughput microbial community analysis on the Illumina HiSeq and MiSeq platforms. ISME J 6:1621–1624.

37. Mikheyev AS, Tin MMY. 2014. A first look at the Oxford Nanopore MinION sequencer. Mol Ecol Resour 14:1097–1102.

38. Seemann T. 2014. Prokka: Rapid prokaryotic genome annotation. Bioinformatics 30:2068–2069.

39. Bortolaia V, Kaas RS, Ruppe E, Roberts MC, Schwarz S, Cattoir V, Philippon A, Allesoe RL, Rebelo AR, Florensa AF, Fagelhauer L, Chakraborty T, Neumann B, Werner G, Bender JK, Stingl K, Nguyen M, Coppens J, Xavier BB, Malhotra-Kumar S, Westh H, Pinholt M, Anjum MF, Duggett NA, Kempf I, Nykäsenoja S, Olkkola S, Wieczorek K, Amaro A, Clemente L, Mossong J, Losch S, Ragimbeau C, Lund O, Aarestrup FM. 2020. ResFinder 4.0 for predictions of phenotypes from genotypes. J Antimicrob Chemother 75:3491–3500.

40. Gupta SK, Padmanabhan BR, Diene SM, Lopez-Rojas R, Kempf M, Landraud L, Rolain J-M. 2014. ARG-ANNOT, a new bioinformatic tool to discover antibiotic resistance genes in bacterial genomes. Antimicrob Agents Chemother 58:212–20.

41. Alcock BP, Raphenya AR, Lau TTY, Tsang KK, Bouchard M, Edalatmand A, Huynh W, Nguyen AL V., Cheng AA, Liu S, Min SY, Miroshnichenko A, Tran HK, Werfalli RE, Nasir JA, Oloni M, Speicher DJ, Florescu A, Singh B, Faltyn M, Hernandez-Koutoucheva A, Sharma AN, Bordeleau E, Pawlowski AC, Zubyk HL, Dooley D, Griffiths E, Maguire F, Winsor GL, Beiko RG, Brinkman FSL, Hsiao WWL, Domselaar G V., McArthur AG. 2020. CARD 2020: Antibiotic resistome surveillance with the comprehensive antibiotic resistance database. Nucleic Acids Res 48:D517–D525.

42. Carattoli A, Hasman H. 2020. PlasmidFinder and In Silico pMLST: Identification and Typing of Plasmid Replicons in Whole-Genome Sequencing (WGS), p. 285–294. In Methods in Molecular Biology. Humana Press Inc.

43. Stothard P, Grant JR, Van Domselaar G. 2018. Visualizing and comparing circular genomes using the CGView family of tools. Brief Bioinform 20:1576–1582.

44. Shannon P, Markiel A, Ozier O, Baliga NS, Wang JT, Ramage D, Amin N, Schwikowski B, Ideker T. 2003. Cytoscape: A software Environment for integrated models of biomolecular interaction networks. Genome Res 13:2498–2504.

45. Rebelo AR, Bortolaia V, Kjeldgaard JS, Pedersen SK, Leekitcharoenphon P, Hansen IM, Guerra B, Malorny B, Borowiak M, Hammerl JA. 2018. Multiplex PCR for detection of plasmid-mediated colistin resistance determinants, mcr-1, mcr-2, mcr-3, mcr-4 and mcr-5 for surveillance purposes. Eurosurveillance23.

46. Nabti LZ, Sahli F, Ngaiganam EP, Radji N, Mezaghcha W, Lupande-Mwenebitu D, Baron SA, Rolain JM, Diene SM. 2020. Development of real-time PCR assay allowed describing the first clinical Klebsiella pneumoniae isolate harboring plasmid-mediated colistin resistance mcr-8 gene in Algeria. J Glob Antimicrob Resist 20:266–271.

